# Evolution of Hierarchical Phase-Contrast Tomography on the European Synchrotron beamlines BM05 and BM18: a whole adult human brain imaging case study

**DOI:** 10.64898/2026.01.15.699653

**Authors:** Hector Dejea, Joseph Brunet, Theresa Urban, Camille Berruyer, Alessandro Mirone, Christoph Jarnias, Alexandre Bellier, Claire Walsh, Peter D. Lee, Paul Tafforeau

## Abstract

Hierarchical Phase-Contrast Tomography (HiP-CT) was recently developed to enable the ex-vivo imaging of human organs at multiple scales from whole organ down to cellular level. Using whole adult human brain imaging as a case study, this manuscript shows the evolution of this technique from its initial development at the BM05 beamline to its transition and current status at BM18. Thanks to the higher coherence, larger beam size, higher energies and larger propagation distances available at BM18 and due to the European Synchrotron’s Extremely Brilliant Source upgrade (ESRF-EBS), this transition resulted in significantly improved data quality, resolution, sensitivity and speed. More recently, the implementation of a new generation of larger sCMOS cameras, helical scanning (including dedicated reconstruction algorithm developments), binning at the chip and projections levels, and the design of high-efficiency optics allowed to progressively improve the trade-off between dose and image quality, while simultaneously reducing scanning times. All these acquisition schemes present the current status of full organ imaging using HiP-CT and represent the constant efforts for the improvement of the technique towards the investigation of human organs in health, disease and aging.

**Synopsis:** This manuscript shows the evolution of Hierarchical Phase-Contrast Tomography (HiP-CT) from its origins at the BM05 beamline to its transition to BM18 (ESRF-EBS). The novel hardware and scanning approach developments resulted in significantly improved data quality, resolution, sensitivity and speed. The case of whole adult brain imaging is presented to demonstrate the current possibilities of full organ imaging with local micron resolution using HiP-CT.

## 1. Introduction

With the wide availability of 3rd generation synchrotron facilities during the last decades, propagation-based X-ray phase-contrast tomography has become a staple methodology for non-destructive, three-dimensional (3D), high-resolution studies in a broad variety of fields, such as biomedicine, palaeontology, earth and material sciences, among others.

Specially for biomedical applications, X-ray phase-contrast imaging has been able to overcome the limitation of resolving low absorption contrast in soft-tissues by exploiting the differences in refractive index, which are orders of magnitude more sensitive in the typical energy range used for X-ray imaging (Lewis, 2004). Propagation-based phase-contrast imaging has been applied to study a multitude of organs down to the cellular level: lungs (Lovric *et al*., 2013; Norvik *et al*., 2020; Borisova *et al*., 2021; Fardin *et al*., 2021); heart (Mirea *et al*., 2015; Gonzalez-Tendero *et al*., 2017; Dejea *et al*., 2019; Wang *et al*., 2019; Reichardt *et al*., 2020; Planinc *et al*., 2021); brain (Beltran *et al*., 2011; Massimi *et al*., 2019; Longo *et al*., 2021; Bosch *et al*., 2022); musculoskeletal tissues (Cooper *et al*., 2011; Disney *et al*., 2019; Pierantoni *et al*., 2021; Madi *et al*., 2020; Dejea *et al*., 2024), pancreas (Tapfer *et al*., 2013; Frohn *et al*., 2020; Pinkert-Leetsch *et al*., 2023); and many others. However, these studies have been limited to small animal (usually rodent) organs or tissue biopsies of human or large animals due to the technical specifications of available beamlines, primarily small beam-sizes (generally < 50 x 10 mm^2^) and limited propagation distances, without the level of coherence required to avoid image blurring at long propagation distances. Slightly larger samples (up to a few centimetres) have been imaged using other phase-contrast techniques less demanding in terms of coherence and propagation, such as grating interferometry, diffraction enhanced imaging, speckle tracking and edge illumination (Keyriläinen *et al*., 2005; Beltran *et al*., 2011; Bravin *et al*., 2012; Diemoz *et al*., 2016; Kaneko *et al*., 2017). However, these methods are dose-inefficient compared to propagation-based techniques, require more cumbersome setups, and have yet to image intact adult organs at micron resolution.

With the recent, current and near-future upgrade of light sources around the world into 4th generation synchrotrons, such as the Extremely Brilliant Source upgrade at the European Synchrotron (ESRF-EBS), highly coherent (100x higher than former ESRF (Pacchioni, 2019)) high-flux beams have enabled high-energy full-field tomography beamlines to achieve propagation distances reaching in some cases the near-field limit for single-distance propagation without being limited by the coherence, so that the capability to resolve small density differences has been dramatically increased. At the ESRF-EBS this has been especially significant for previous bending magnet beamlines, which have been substituted by short two or three-pole wigglers and can now take full advantage of the upgrade benefits.

Taking advantage of these developments, Hierarchical Phase-Contrast Tomography (HiP-CT) (Walsh et al., 2021) was developed to enable the imaging of adult human organs at multiple scales from whole organ down to near cellular level. HiP-CT has already been successfully applied to the study of several adult human organs in health and disease, such as lung (Ackermann, Kamp *et al*., 2022; Ackermann, Tafforeau *et al*., 2022; Verleden *et al*., 2024), heart (Brunet *et al*., 2024), placenta (Reichmann *et al*., 2024) or temporal bone (Schaeper *et al*., 2025).

HiP-CT was initially developed at the ‘’X-ray Imaging and Optics Test” beamline BM05 (ESRF-EBS), which is limited to a maximum propagation distance of 3.5 m and a beam size of 50 x 4 mm^2^ at the sample position (52 m). However, the recent beamline for “Hierarchical Tomography” BM18 (ESRF-EBS) (Lang *et al*., 2023; Cianciosi *et al*., 2023; Vijayakumar *et al*., 2024) takes full advantage of the enhanced EBS brilliance properties by allowing up to 36 m of propagation in a 220 m long beamline, with a filtered polychromatic beam (38 – 300 keV) of 300 x 17 mm^2^ at the sample position (178 m). See Table 1 for comparative technical specifications of both beamlines.

**Table 1.**
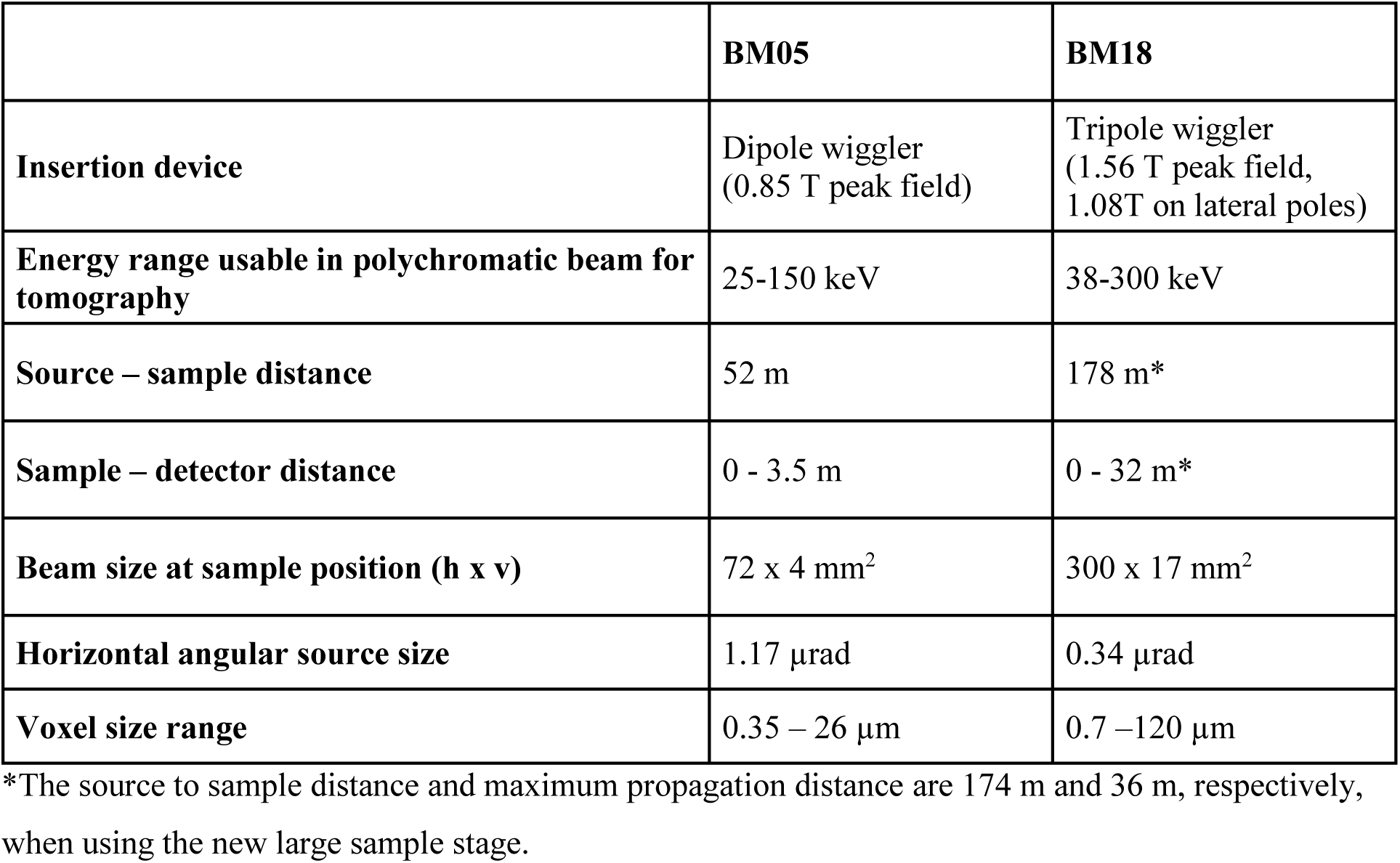
Technical specifications of BM05 and BM18 beamlines (ESRF-EBS).

In this manuscript, we aim to report on developments enabling the evolution of HiP-CT data quality, resolution and sensitivity. These have been enabled by the transition from BM05 to BM18, as well as the incorporation of new hardware, alternative acquisition approaches, and software developments. This is demonstrated by using the case study of whole adult human brain imaging.

## 2. Material and Methods

### 2.1. Organ and donor description

A whole control human brain was obtained from a body donation to the Department of Anatomy (LADAF). The donor LADAF-2021-17 was a 63-year-old, 178 cm, 60 kg male with pancreatic cancer. During autopsy, stomach hypertrophy and aspiration of gastric contents were noted.

Body donation is based on a voluntary act of free consent by the donors antemortem. The dissection was performed respecting the memory of the deceased and following current French legislation for body donation. The post-mortem study was conducted according to the Quality Appraisal for Cadaveric Studies scale recommendations (Wilke et al., 2015).

### 2.2. Organ preparation

After death, the donor body was embalmed by injecting formalin-based embalming solution into the right carotid artery. The body was then stored at 3.6 °C for a few days. The dissection, including the extraction of the brain, was performed at the LADAF. The brain was then post-fixed in 4% neutral buffered formalin at 3.6 °C for at least 4 days. Following the sample preparation protocol described by Brunet et al. (Brunet et al., 2023), the brain underwent partial dehydration using increasing ethanol concentration baths until 70% ethanol was reached (4 days per bath at 50%, 60%, 70% and 70% ethanol concentration). The brain was stabilised in equilibrated crushed agar-ethanol gel with the frontal cortex facing downwards and degassed using thermal cycling (Brunet *et al*., 2023). The container was finally sealed and stored at room temperature until scanning (Supplementary Figure S1).

### 2.3. Imaging protocols

To assess the evolution in HiP-CT data quality and compare each beamline, the brain was initially imaged at BM05 using a 25.25 µm voxel size configuration. It was later imaged at BM18 using two configurations at 42.4 µm and 23.42 µm voxel size. Following the technical evolution of the BM18 beamline described below, the same brain was imaged again with consecutive developments at 19.28 µm/voxel, 20.17 µm/voxel and 14.29 µm/voxel. The technical specifications of the beamlines are summarised in Table 1, a detailed list of all setups and acquisition parameters can be found in Supplementary Excel and a timeline of development can be found in Supplementary Figure S2.

#### 2.3.1. BM05 – Initial development

The brain was initially imaged using HiP-CT at the BM05 beamline at the European Synchrotron (ESRF-EBS, Grenoble, France). This dataset can be found at https://doi.org/10.15151/ESRF-DC-1773964937.

The scanning protocol consisted of using a quasi-parallel polychromatic beam attenuated by 0.23 mm of copper and 0.68 mm of molybdenum. Propagation-based full-field tomography of the whole brain was performed in quarter-acquisition mode (a central half-acquisition plus a surrounding “annular” half-acquisition”) at a pixel size of 25.25 µm with a propagation distance of 3.5 m, 9900 projections per scan and 36 ms exposure time (accumulation of 6 times 6 ms). A total of 75 pairs of scans were required to cover the whole brain. X-rays were converted to visible light by a LuAG:Ce 1000 µm scintillator (Crytur, Czech Republic), demagnified by a Dzoom optic (x0.25) and detected by a PCO Edge 4.2 CLHS detector (2048×2048 pixels of 6.5µm, PCO imaging, Germany). This configuration resulted in an average detected energy of 97 keV after absorption by the sample. The total acquisition time was 18.3 h.

#### 2.3.2. BM18 – Improvement of data quality due to long propagation phase-contrast

The brain was also imaged using HiP-CT in five different configurations at the BM18 beamline at the European Synchrotron (ESRF-EBS, Grenoble, France) from 2021-2025 as the technique evolved.

The scanning protocols included the use of a filtered quasi-parallel polychromatic beam. For the first configuration, performed overnight on the first day of tomography on the new BM18 beamline, propagation-based full-field tomography of the whole brain was achieved with the use of a 1.3 mm of molybdenum attenuator, in half-acquisition mode at a pixel size of 42.4 µm with a propagation distance of 38 m, 6000 projections per scan and 105 ms exposure time (accumulation of 7 times 15 ms). A total of 24 vertical scans were required to cover the full volume of the brain. X-rays were converted to visible light by a LuAG:Ce 2000 µm scintillator (Crytur, Czech Republic), demagnified by the LAFIP2 optic (tandem combining a 400mm custom objective with a canon 50 mm one, leading to a x0.125 magnification) and detected by a PCO Edge 4.2 CLHS detector (PCO imaging, Germany). This configuration resulted in an average detected energy of 122 keV after absorption by the sample. The total acquisition time was 5.7 h. This dataset can be found at https://doi.org/10.15151/ESRF-DC-2313098569.

For the second configuration, performed in February 2022 right after the implementation of the Dzoom optic on BM18 (similar to that used on BM05), the organ jar was immersed in 70% ethanol (5 mm thickness) in a surrounding rig tube to guarantee a cylindrical shape and be able to include the complete jar in the field of view without the artefacts coming from the interface with air. Propagation-based full-field tomography was performed with 0.61 mm of molybdenum, in quarter-acquisition mode at a pixel size of 23.42 µm with a propagation distance of 31 m, 9990 projections per scan and 45 ms exposure time (accumulation of 3 times 15 ms). A total of 33 vertical scans were required to cover the whole brain. X-rays were converted to visible light by a LuAG:Ce 2000 µm scintillator (Crytur, Czech Republic), demagnified by the Dzoom optic and detected by a PCO Edge 4.2 CLHS detector (PCO imaging, Germany). This configuration resulted in an average detected energy of 110 keV after absorption by the sample. The total acquisition time was 8.9 h. Regarding the detector configuration, except the thickness of the scintillator that was twice higher on BM18 than on BM05, the two detector systems can be considered as identical. This dataset can be found at https://doi.org/10.15151/ESRF-DC-1773964905.

#### 2.3.3. BM18 – Improving dose efficiency

In the configuration for dose efficiency maximisation in whole organ imaging, the key element was the use of binning at the detector level for an increased detection efficiency (4 pixels in the place of 1), thus reducing the dose required for imaging. Moreover, the addition of large field detectors (Iris 15 Scientific CMOS, Teledyne Photometrics, USA with 5056×2960 pixels of 4.25 µm) allowed to still cover the complete brain in quarter-acquisition mode.

The organ jar was also immersed in 70% ethanol (5 mm thickness). Propagation-based full-field tomography was performed with 10 mm of sapphire, 0.3 mm of silver and 30 mm of glassy carbon, in quarter-acquisition mode at a pixel size of 19.28 µm (9.64 µm in bin 2) and with a propagation distance of 20 m, 12,000 projections per scan and 36 ms exposure time (accumulation of 3 times 12 ms). 24 vertical scans were required to cover the whole brain. X-rays were converted to visible light by a GAGG 1000 µm scintillator with reflective layer (Crytur, Czech Republic), demagnified by a Dzoom optic and detected by an Iris 15 Scientific CMOS detector (Teledyne Photometrics, USA). This configuration resulted in an average detected energy of 103 keV after absorption by the sample. The total acquisition time was 6.8 h. This dataset can be found at https://doi.esrf.fr/10.15151/ESRF-DC-2313101083.

#### 2.3.4. BM18 – Improving acquisition speed

In this configuration, the key additions were the use of helical scanning, which gets rid of overheads in individual scans and improves concatenation artefacts, as well as the use of bin mean (binning performed at the projection level using averaging, rather than on-chip pixel summation).

The organ jar was also immersed in 70% ethanol (5 mm thickness). Propagation-based full-field tomography was performed with 10 mm of sapphire, 0.4 mm of silver and 35 mm of glassy carbon, in quarter-acquisition mode at a pixel size of 20.17 µm (10.08 µm in bin mean 2) and with a propagation distance of 11 m, 12,000 projections per scan and 12 ms exposure time. 35 vertical turns were required to cover the whole brain. X-rays were converted to visible light by a GAGG 1000 µm scintillator with reflective layer (Crytur, Czech Republic), demagnified by a Dzoom optic and detected by an Iris 15 Scientific CMOS detector (Teledyne Photometrics, USA). This configuration resulted in an average detected energy of 113 keV after absorption by the sample. The total acquisition time was 3.2 h. This dataset can be found at https://doi.org/10.15151/ESRF-DC-2313101075.

#### 2.3.5. BM18 – Best configuration with fixed x0.5 magnification optics

This configuration takes advantage of the latest addition of optics at BM18, a fixed magnification tandem optic at x0.5 (Mamiya 200mm f/d 2.8 / Otus 100mm f/s 1.4), which improves significantly the optical efficiency compared to the previously used Dzoom. Based on optical calculations, the gain is approximately a factor 22 for identical scintillator configurations, while experimental measurements indicate an overall signal gain of about 8 to 10 when combined with thinner scintillators (250um GAGG+ with reflective layer), depending on beam energy.

The organ jar was also immersed in 70% ethanol (5 mm thickness). Propagation-based full-field tomography was performed with 5 mm of sapphire and 0.5 mm of silver, in quarter-acquisition mode at a pixel size of 14.79 µm (7.39 µm in bin mean 2) and with a propagation distance of 20 m, 15,000 projections per scan and 18 ms exposure time. 39 helical turns were required to cover the whole brain. X-rays were converted to visible light by a GAGG 250 µm scintillator with reflective layer (Crytur, Czech Republic), demagnified by a fixed x0.5 magnification optic and detected by an Iris 15 Scientific CMOS detector (Teledyne Photometrics, USA). This configuration resulted in an average detected energy of 102 keV after absorption by the sample. The total acquisition time was 6.5 h. This dataset can be found at https://doi.org/10.15151/ESRF-DC-2313101091.

### 2.4. Tomographic reconstruction and data visualization

The initial 3 scans were reconstructed following the pipeline described in detail by (Xian et al., 2022). Briefly, the reconstruction consisted of a filtered back-projection algorithm in combination with the single-distance phase retrieval algorithm by Paganin et al. (Paganin et al., 2002) and 2D unsharp filtering using PyHST2 (Mirone et al., 2014). For the two first datasets (BM05 at 25.25 µm and BM18 at 42.4 µm voxel size), the reconstructed volumes were vertically concatenated to obtain the full brain volume. For all subsequent acquisitions, the vertical concatenation was instead performed at the projection level prior to reconstruction. Ring artefact correction was then performed on the reconstructed slices using an algorithm based on (Lyckegaard et al., 2011).

The 2 final helical scans were reconstructed using the ESRF tomography processing software Nabu together with the night rail framework (Mirone, 2025). The presented data have been registered and visualized using VGSTUDIO MAX 2024.1.

All the reconstruction are available free-access at the Human Organ Atlas portal (https://human-organ-atlas.esrf.fr) (Walsh *et al*., 2025).

## 3. Results

Figures 1, 2 and 3 illustrate the improvement in HiP-CT data quality by comparing a scan of a whole adult human brain (LADAF-2021-17) at BM05 at 25.25 µm/voxel to measurements of the same brain at BM18 with 42.4 µm/voxel, 23.42 µm/voxel, 19.28 µm/voxel, 20.17 µm/voxel and 14.79 µm/voxel – each of these measurements representing an evolution aspect of the technique. The figures present data in the coronal, sagittal and transversal planes, respectively.

**Figure 1.**
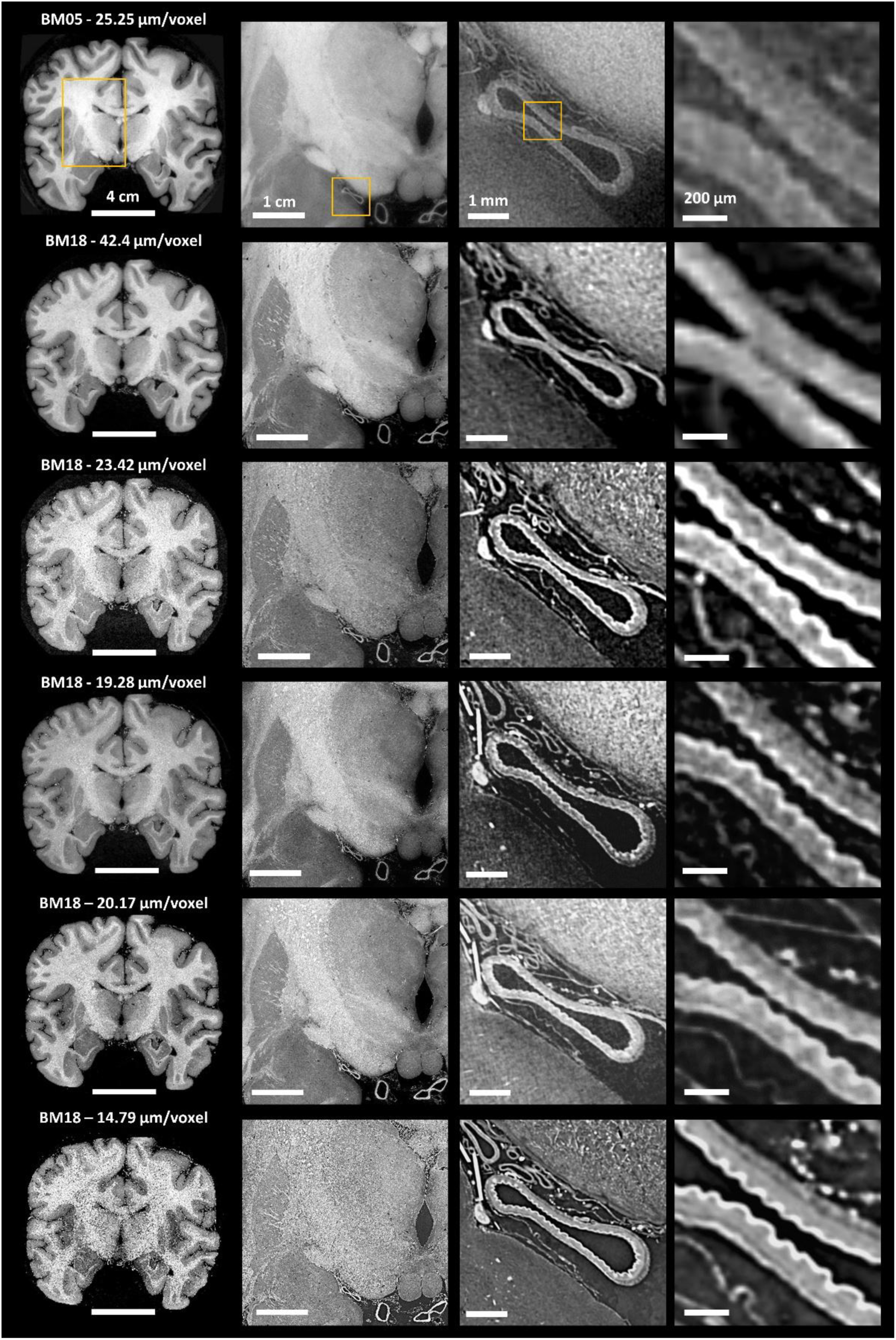
Comparison between BM05 and BM18 acquisitions using an illustrative coronal slice at the same location of the brain. From left to right column: complete slice, first, second and third digital zoom levels in one of the main arteries. Zoom locations are depicted in yellow in the top row and remain equal across acquisition types.

**Figure 2.**
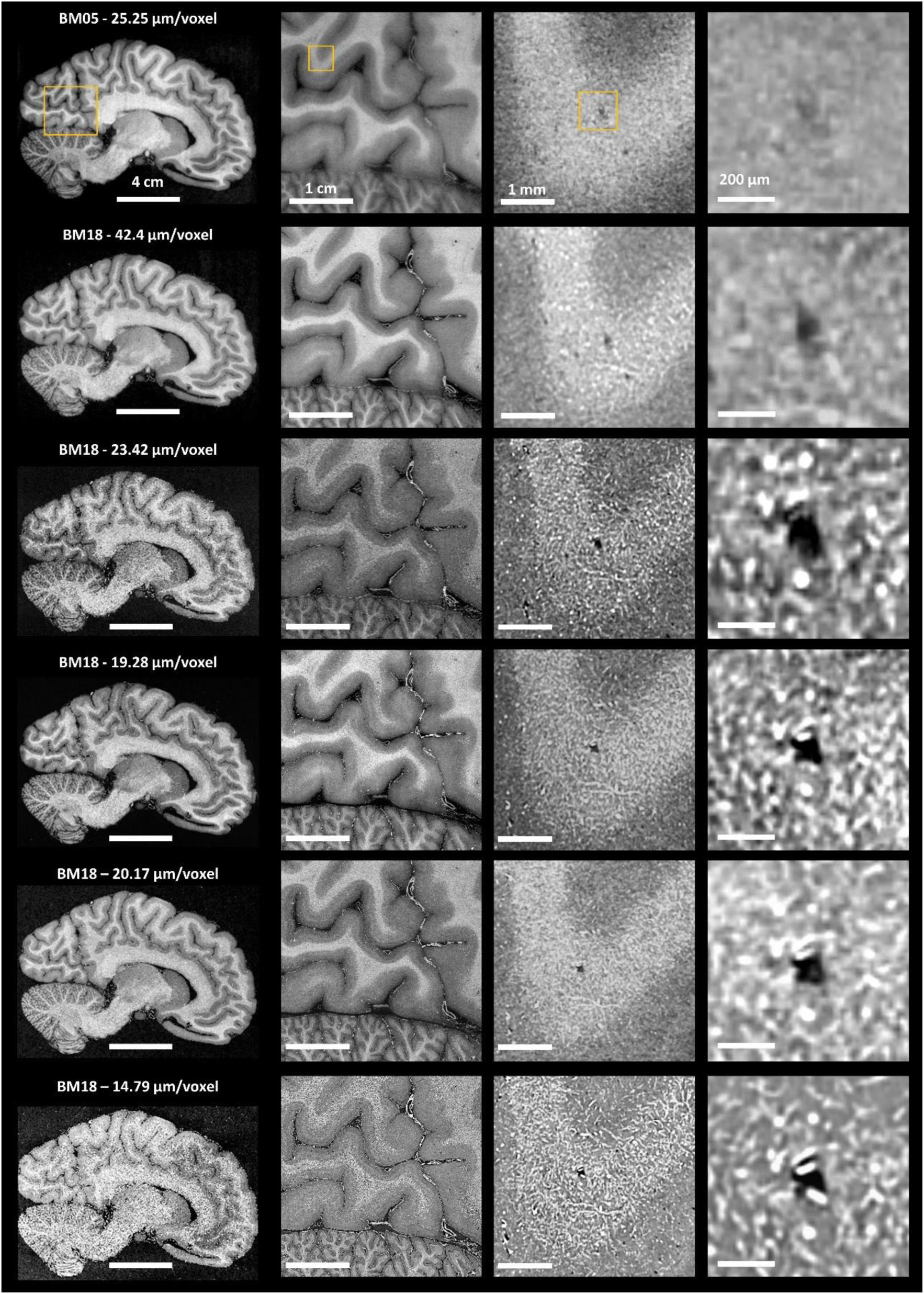
Comparison between BM05 and BM18 acquisitions using an illustrative sagittal slice at the same location of the brain. From left to right column: complete slice, first, second and third digital zoom level in the cortex containing both grey and white matter. Zoom locations are depicted in yellow in the top row and remain equal across acquisition types.

**Figure 3.**
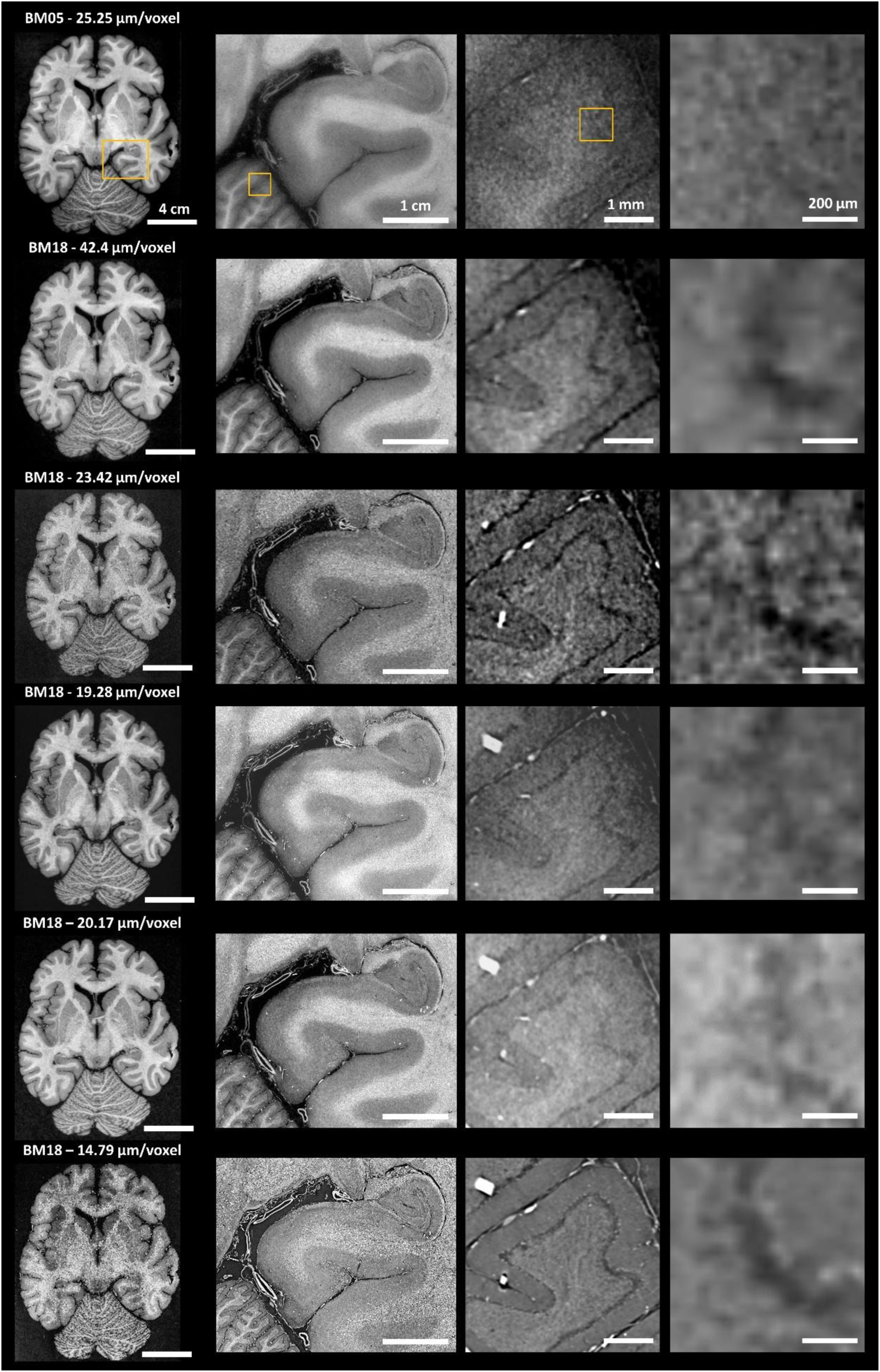
Comparison between BM05 and BM18 acquisitions using an illustrative transversal slice at the same location of the brain. From left to right column: complete slice, first, second and third digital zoom level of the cerebellum. Zoom locations are depicted in yellow in the top row and remain equal across acquisition types.

The BM05 data (25.25 µm/voxel) shows good contrast between white and grey matter, but is unable to resolve structures within the tissue other than the ventricles and very large vessels. This is due to the very short propagation distance, which does not allow small structures with weak phase shifts to generate sufficient phase fringes, thus losing sensitivity. The improvement in data quality is already very clear for BM18 data at 42.4 µm/voxel, which is able to retain high contrast while improving the visualization of internal structures and tissue boundaries. Using a comparable pixel size of 23.42 µm/voxel, the improvement to sensitivity is dramatic, enabling even different arterial sub-layers (intima, media, adventitia) in large vessels to be resolved (Figure 1).

The incorporation of large chip detectors at BM18 (Iris15, 5056 x 2960 pixels, 4.25 µm/pixel) allowed covering much larger field of views at smaller voxel size. Initially in combination with chip binning, and later with helical scanning and binning at the projection level, the dose deposited could be reduced while keeping imaged quality and resolving very similar structures (Figures 1-3, 19.28 µm/pixel and 20.17 µm/pixel).

Finally, the development of a fixed magnification x0.5 clearly shows a great improvement in image quality while again reducing the dose required. This can be clearly noted by the sharpness of structures in vasculature, grey-white matter and cerebellum (Figures 1-3, 14.79 µm/pixel).

## 4. Discussion

This manuscript presents the evolution of the HiP-CT data quality by transitioning from its original development on beamline BM05 to BM18 (ESRF-EBS), which is purposely designed for multi-resolution phase contrast imaging. It also shows the technique’s evolution to lower dose and scanning times thanks to the incorporation of new hardware and software tools, as well as alternative acquisition schemes (Supplementary Figure S2).

The technical specifications of BM05, a relatively short beamline in comparison to BM18 (52 vs 178 m source-sample distance) and limited propagation distance (maximum 3.5 vs 38 m), make it ideal for phase contrast imaging below ∼4 µm. Assuming an energy of 100 keV, the near-field limit is constrained by the hutch size at 3.3 µm/voxel, and if the hutch was larger, geometrical blurring will be already dominating at a propagation distance of 5 m for voxel sizes larger than 4 µm (Figure 4). This is not the case for BM18, which allows to approach propagation distances very close to the near-field limit without reaching geometrical blurring, so that the limitation becomes the size of the experimental hutch rather than the horizontal angular source size (Figure 4). Assuming an energy of 100 keV, a propagation distance of 59.4 m would be required for geometrical blurring to dominate over the near-field limit for voxel sizes above 13.5 µm, a regime that remains inaccessible within the current hutch dimensions. See Supplementary Information and Supplementary Figure S3 for more details on near-field limit and geometrical blur calculations.

**Figure 4.**
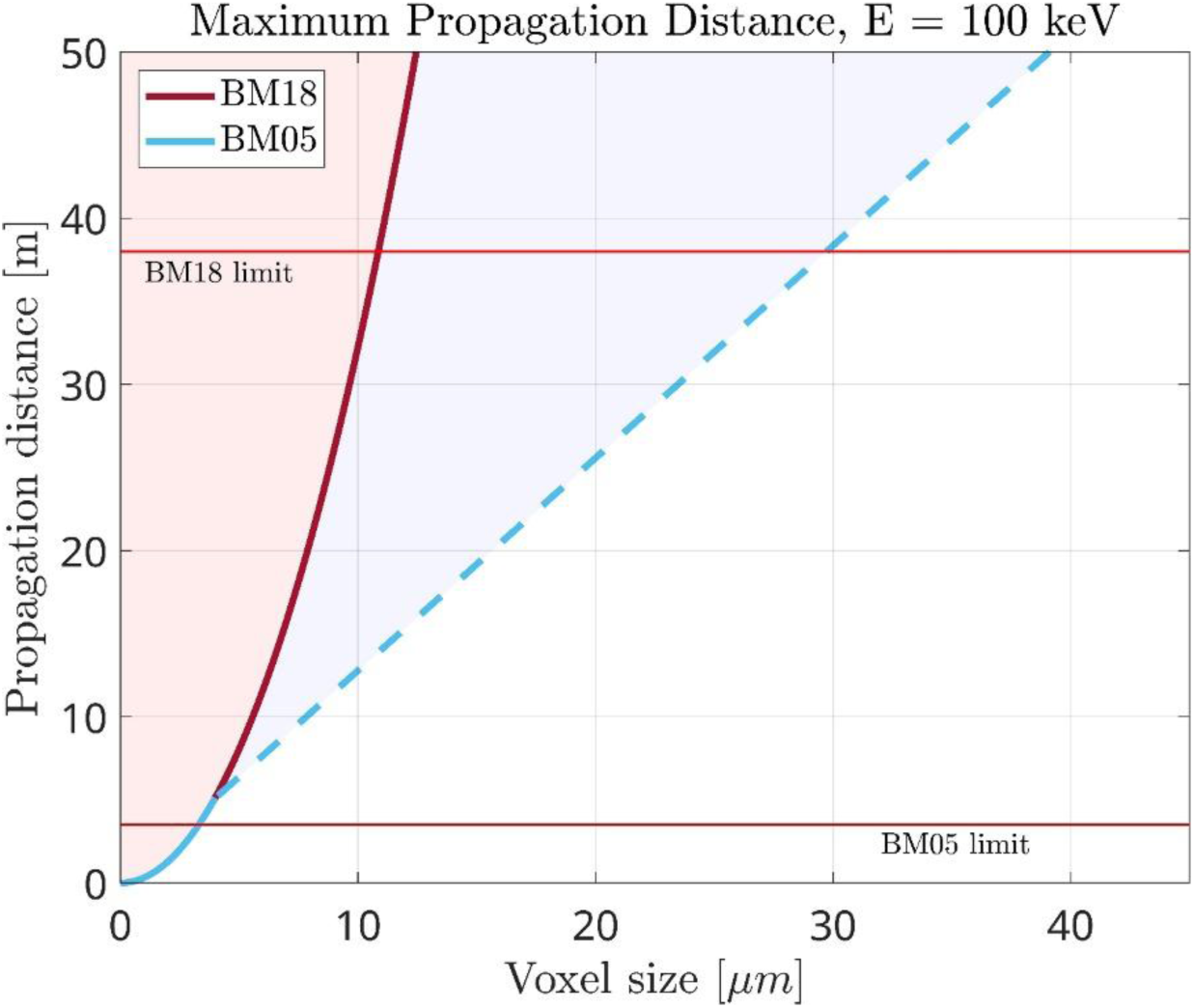
Maximum propagation distance achievable at beamlines BM18 and BM05 for an energy of 100 keV, as constrained by either near-field limit or geometrical blur (independent of energy). The horizontal red lines indicate the maximum propagation distance achievable by hutch size. The dashed curve indicates the region at which geometrical blur takes over the near-field limit, which does not happen in BM18 until much further propagation distances than plotted. Forbidden zones are shadowed with the corresponding beamline color.

As a consequence, the specifications of BM18 make it the best currently existing beamline in the world to perform HiP-CT on samples as large as or even larger than adult human organs, which is clearly visible by the comparison in Figures 1, 2, 3. The larger propagation distance at such level of coherence is able to take full advantage of the phase contrast effect and amplify smaller structures, so that the overall data quality and sensitivity are improved for similar acquisition in both beamlines.

While BM18 is proven here to show a remarkable superiority in whole organ imaging, BM05 remains a very powerful instrument for sub-micron voxel size imaging thanks to its higher power density (at < 140 keV) and should always be considered for micrometre-resolution studies, as long as the propagation distance and smaller beam size do not become limitations.

After the transition from BM05 to BM18, the development of the technique has focused on the optimization of image quality with minimum dose deposition and scanning times. Large chip detectors with smaller pixel size (Iris15) were incorporated with the goal to improve resolution and data quality while reducing the dose and data size, which could be achieved by working in binning at the chip level. This allowed to maintain image quality while reducing the dose by ∼75 % (Supplementary Excel and Figure 5). However, this acquisition protocol still required the use of accumulation to reduce the noise level, which could be avoided by the use of binning by average at the projection level, instead of at the chip level (bin mean). This means that with a single shot, it is possible to achieve a dynamic level corresponding to an accumulation of 4 (coming from the 4 averaged pixels) without saturating the 32 bits dynamic scale of the pictures. Additionally, in the meantime, the possibility to perform helical scanning was developed at BM18, thus allowing to reduce scanning times by avoiding overheads in each stacked scan and obtaining an equivalent of accumulation 2 by doing a 50 % overlap between helical turns. These advancements resulted in an approximately 50% reduction in both total scanning time and integrated absorbed dose, as summarised in Figure 5.

**Figure 5.**
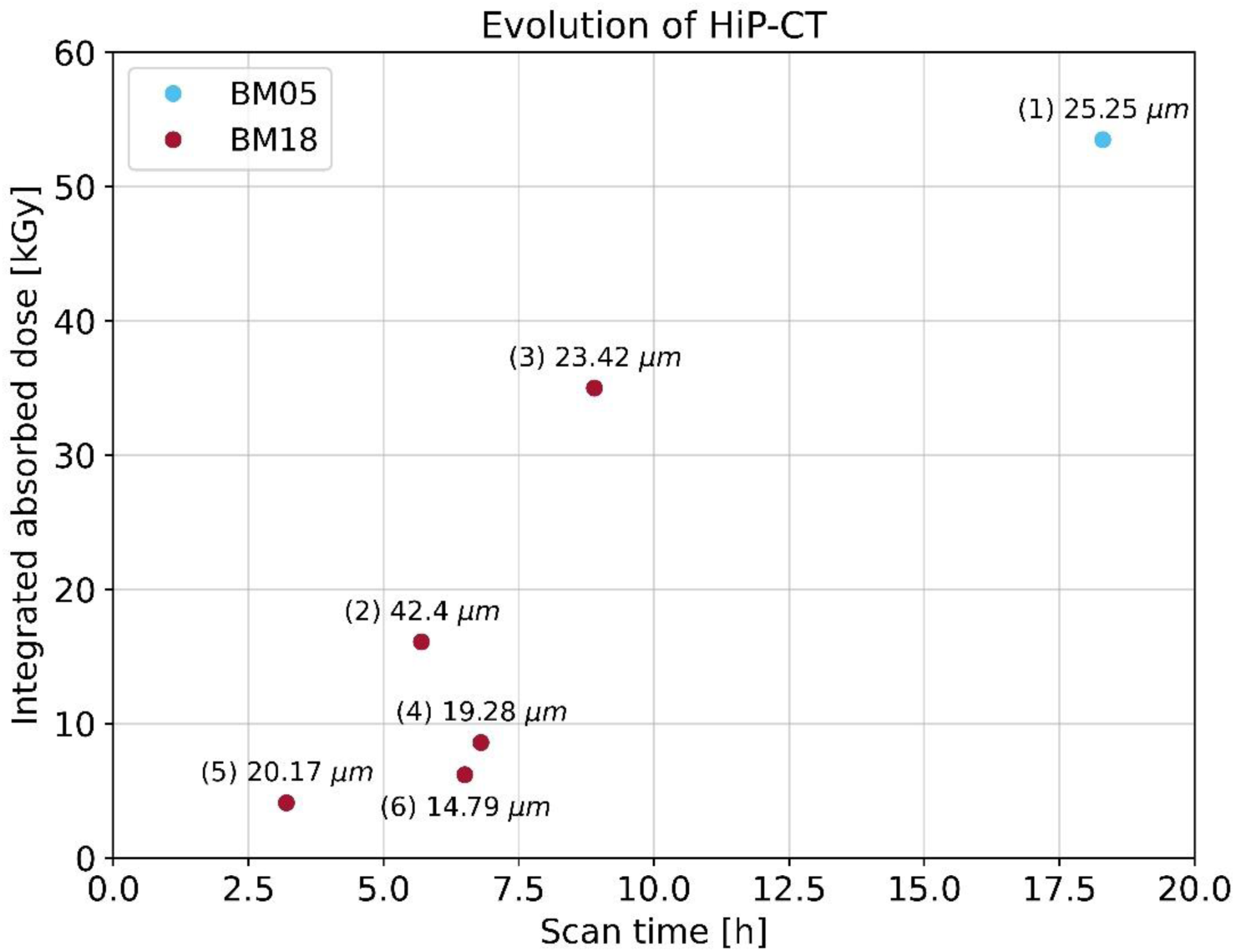
Relationship between integrated absorbed dose and total scanning time for the presented setups at beamlines BM18 and BM05. Each data point is marked by the setup number in order of acquisition and the corresponding pixel size.

The final step corresponds to a major improvement in image quality and sensitivity enabled by the introduction of a fixed magnification x0.5. This optic is approximately 32 times more efficient than the previously used DZoom in term of photon collection, enabling substantially higher sensitivity and improved image quality at smaller voxel sizes. In practice, this gain is primarily invested in improving spatial resolution and contrast, and acquisition speed rather than reducing dose, allowing a reduction of the voxel size from approximately 20 µm to 15 µm while maintaining comparable dose levels. This improvement arises primarily from the much larger numerical aperture of the fixed x0.5 optic compared to the DZoom system (NA ≈ 0.2 versus 0.042), leading to a strongly enhanced light collection efficiency. Additional gains result from improved optical transmission and the use of more efficient scintillators (GAGG+ instead of LuAG:Ce). The higher numerical aperture also improves the diffraction-limited resolution, implying the use of thinner scintillators that brings higher sensitivity and image sharpness. As a result, this configuration becomes thus a game changer for the scanning of organ overviews with maximum efficiency.

This manuscript has shown the developments, current status and possibilities of whole organ imaging with HiP-CT after its successful transition from BM05 to BM18, using the same adult human brain as case study. This evolution in configurations presents a reduction in integrated dose values (Figure 5), which is very beneficial to reduce the risk of bubble formation (Xian *et al*., 2024) and damage at the molecular level for further downstream “omics” analyses. To our knowledge, no other imaging technique is able to provide whole adult human organ data in 3D, non-destructively, and at such spatial resolution and speed. That said, cumulative data size (∼1 TB per organ) and processing or analysis time, which is currently highly manual, remain a bottleneck. To tackle this, more efforts need to be put on the development of artificial intelligence methods and tools that can speed up the process while providing reliable quantification (Greenspan *et al*., 2016; Hesamian *et al*., 2019; Yagis *et al*., 2024).

Currently, HiP-CT is fully operational at both BM05 and BM18 beamlines of the ESRF-EBS. In the near future, on-going hardware developments, such as the recently commissioned big sample stage (max 300 kg sample, 2.3 m vertical translation, 1.2 m sample diameter) in combination with existing optics that can take the full 30 cm of beam, will allow imaging of larger multi-organs complexes. In addition, the recent installation of a PCO Edge 10 bi CLHS detector (4416×2368 pixels of 4.6µm, PCO imaging, Germany) detector coupled to the fixed magnification x0.5 optics enables data acquisition with comparable image quality while reducing scan times by approximately a factor of three for the same dose, significantly improving throughput for large samples. In parallel, the later addition of the next generation “Large Area Detector” (LAD) from Fraunhofer IIS, with a lateral size of 16,000 pixels coming from the combination of 9 sCMOS chips, would enable the possibility to image full ex-vivo human bodies at 20 µm/voxel. In addition, the upgrades of other light sources to 4^th^ generation, such as APS, SLS, and DLS, or the construction of new beamlines such as the Biomedical Imaging Long Beamline project in South Korea, open the possibility of translating this technique for large organs to other facilities.

## Supporting information

Supplementary Excel: Detailed list of all setups and acquisition parameters

## Acknowledgements

We sincerely thank the donors and their families for their invaluable contributions. Results from such research can potentially increase humanity’s overall knowledge and improve patient care. These donors and their families therefore deserve our highest gratitude. We thank Élodie Boller, Clémence Muzelle, Roberto Homs, Filippo Cianciosi, and Philippe Carceller for their contributions to setup development and improvements. We further thank David Stansby and Guillaume Geisne for their support with data management and uploading. This project has been made possible in part by CZI grant DAF2020-225394 and grant DOI https://doi.org/10.37921/331542rbsqvn from the Chan Zuckerberg Initiative DAF, an advised fund of Silicon Valley Community Foundation (funder DOI 10.13039/100014989), the ESRF funding proposals md1252, md1290 and md1389, the Royal Academy of Engineering (CiET1819/10). PDL, CLW and JB gratefully acknowledge funding from the MRC (MR/R025673/1). P.D.L. is a CIFAR MacMillan Fellow in the Multiscale Human Program. The authors declare no conflicts of interest.

## Compliance with Ethical Standards

As part of research involving human participants in the context of body donation, the authors declare that they have complied with current French law (funeral and public health legislation) and have respected the ethical principles of respect for the deceased. The research was conducted in accordance with the 1964 Helsinki Declaration. The research was authorised by the Grenoble Alpes University.

## Supporting information

**Figure S1.**
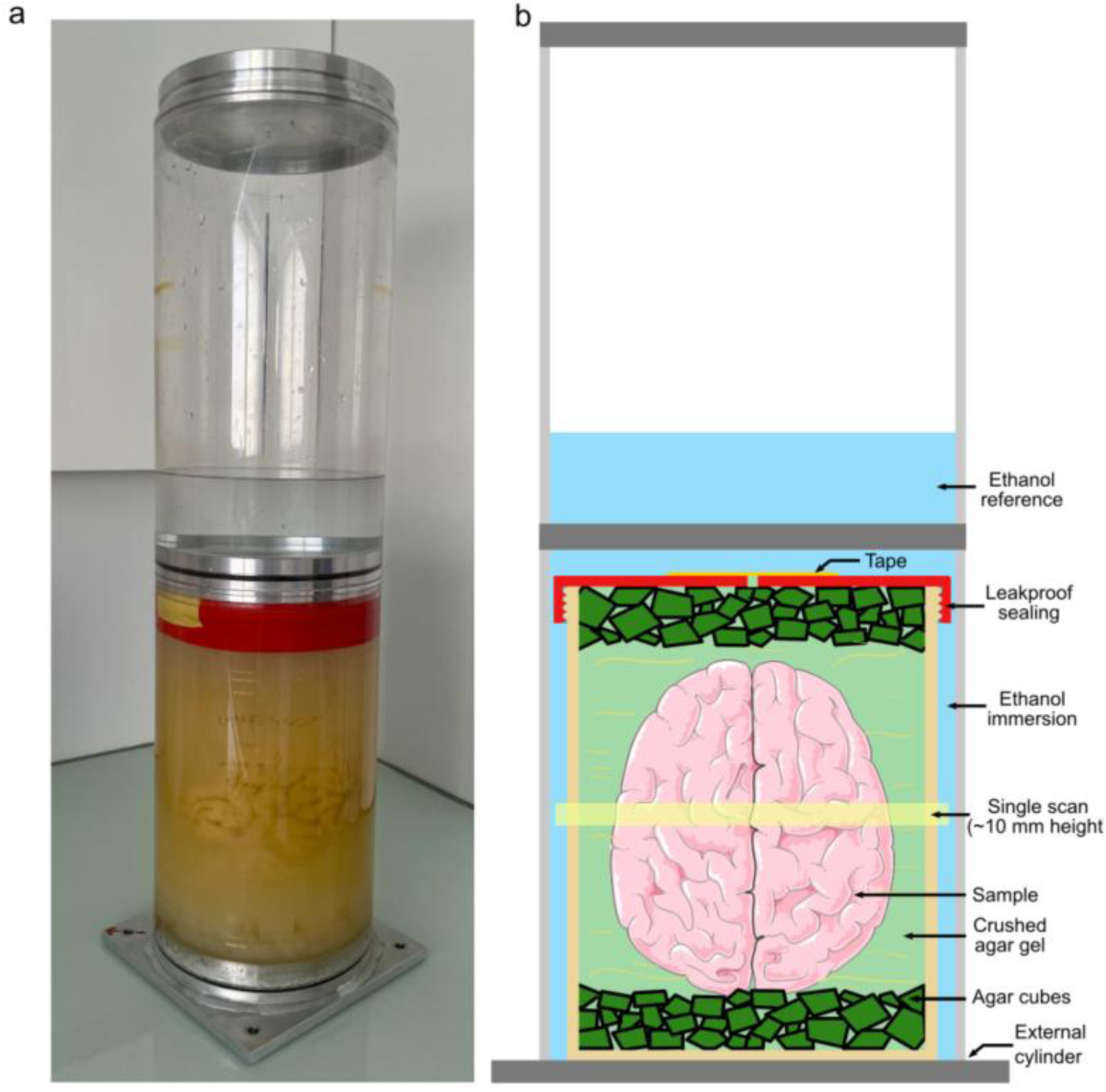
a) Picture and b) schematic of the organ mounting required for HiP-CT.

**Figure S2.**
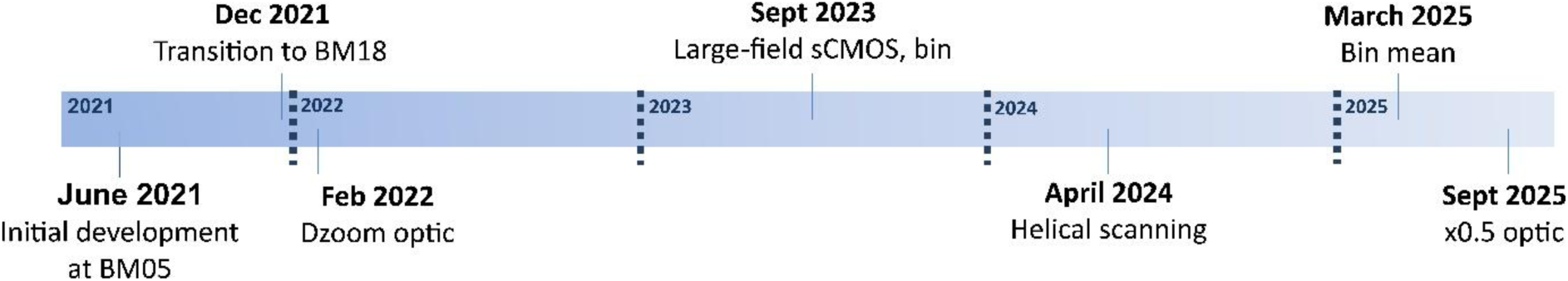
Timeline of development in whole adult human organ imaging at BM05 and BM18.

### S1. Near-field limit and geometrical blur

As described in (Salditt *et al*., 2017), the near-field regime for a sample structure of size *a* measured at a wavelength λ is given by a Fresnel number (*F*) below 1, so that as 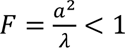. We assumed that at least 2 pixels are required to resolve a structure, so that the critical propagation distance *z_c_* for a specific structure size is given by 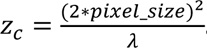 Geometrical blur (*g*) is caused by the finite source size (σ) and the ratio of source-sample distance (*SSD*) and sample-detector distance (*SDD*). For this calculation, we assumed a maximum blur of 1.5 pixels, so that 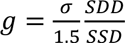.

Supplementary Figure S3 presents the near-field limit and geometrical blur for BM05 and BM18.

**Figure S3.**
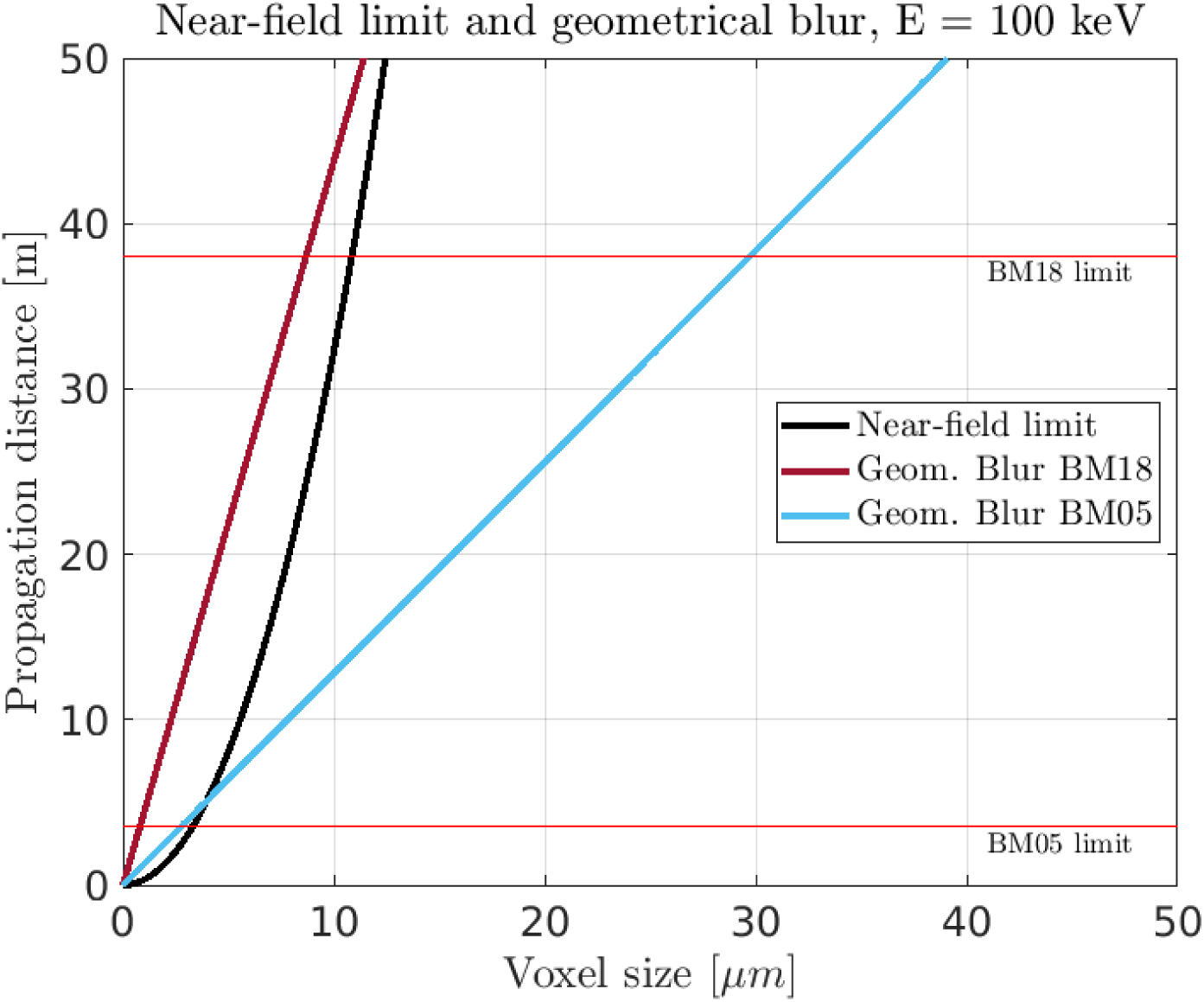
Near-field limit at an energy of 100 keV and geometrical blur curves for BM18 and BM05. Geometrical blur is independent of the energy. The horizontal red lines indicate the maximum propagation distance allowed by the experimental hutches.

## References

Ackermann, M., Kamp, J. C., Werlein, C., Walsh, C. L., Stark, H., Prade, V., Surabattula, R., Wagner, W. L., Disney, C., Bodey, A. J., Illig, T., Leeming, D. J., Karsdal, M. A., Tzankov, A., Boor, P., Kühnel, M. P., Länger, F. P., Verleden, S. E., Kvasnicka, H. M., Kreipe, H. H., Haverich, A., Black, S. M., Walch, A., Tafforeau, P., Lee, P. D., Hoeper, M. M., Welte, T., Seeliger, B., David, S., Schuppan, D., Mentzer, S. J. & Jonigk, D. D. (2022). eBioMedicine 85, 104296.

Ackermann, M., Tafforeau, P., Wagner, W. L., Walsh, C. L., Werlein, C., Kühnel, M. P., Länger, F. P., Disney, C., Bodey, A. J., Bellier, A., Verleden, S. E., Lee, P. D., Mentzer, S. J. & Jonigk, D. D. (2022). Am J Respir Crit Care Med 205, 121–125.

Beltran, M. A., Paganin, D. M., Siu, K. K. W., Fouras, A., Hooper, S. B., Reser, D. H. & Kitchen, M. J. (2011). Phys. Med. Biol. 56, 7353–7369.

Borisova, E., Lovric, G., Miettinen, A., Fardin, L., Bayat, S., Larsson, A., Stampanoni, M., Schittny, J. C. & Schlepütz, C. M. (2021). Histochem Cell Biol 155, 215–226.

Bosch, C., Ackels, T., Pacureanu, A., Zhang, Y., Peddie, C. J., Berning, M., Rzepka, N., Zdora, M.-C., Whiteley, I., Storm, M., Bonnin, A., Rau, C., Margrie, T., Collinson, L. & Schaefer, A. T. (2022). Nat Commun 13, 2923.

Bravin, A., Coan, P. & Suortti, P. (2012). Phys. Med. Biol. 58, R1.

Brunet, J., Cook, A. C., Walsh, C. L., Cranley, J., Tafforeau, P., Engel, K., Arthurs, O., Berruyer, C., Burke O’Leary, E., Bellier, A., Torii, R., Werlein, C., Jonigk, D. D., Ackermann, M., Dollman, K., Lee, P. D. & Atzen, S. (2024). Radiology 312, e232731.

Brunet, J., Walsh, C. L., Wagner, W. L., Bellier, A., Werlein, C., Marussi, S., Jonigk, D. D., Verleden, S. E., Ackermann, M., Lee, P. D. & Tafforeau, P. (2023). Nat Protoc 18, 1441–1461.

Cianciosi, F., Buisson, A.-L., Carceller, P., Tafforeau, P. & van Vaerenbergh, P. (2023). *12th Mechanical Engineering Design of Synchrotron Radiation Equipment and Instrumentation* TUOAM02.

Cooper, D. M. L., Erickson, B., Peele, A. G., Hannah, K., Thomas, C. D. L. & Clement, J. G. (2011). J. Anat. 219, 481–489.

Dejea, H., Garcia-Canadilla, P., Cook, A. C., Guasch, E., Zamora, M., Crispi, F., Stampanoni, M., Bijnens, B. & Bonnin, A. (2019). Sci Rep 9, 6996.

Dejea, H., Pierantoni, M., Orozco, G. A., Bokvist Wrammerfors, E. T., Gstohl, S. J., Schlepütz, C. M. & Isaksson, H. (2024). Adv Sci 2308811.

Diemoz, P. C., Bravin, A., Sztrókay-Gaul, A., Ruat, M., Grandl, S., Mayr, D., Auweter, S., Mittone, A., Brun, E., Ponchut, C., Reiser, M. F., Coan, P. & Olivo, A. (2016). Phys. Med. Biol. 61, 8750–8761.

Disney, C. M., Eckersley, A., McConnell, J. C., Geng, H., Bodey, A. J., Hoyland, J. A., Lee, P. D., Sherratt, M. J. & Bay, B. K. (2019). Acta Biomaterialia 92, 290–304.

Fardin, L., Broche, L., Lovric, G., Mittone, A., Stephanov, O., Larsson, A., Bravin, A. & Bayat, S. (2021). Sci Rep 11, 4236.

Frohn, J., Pinkert-Leetsch, D., Missbach-Güntner, J., Reichardt, M., Osterhoff, M., Alves, F. & Salditt, T. (2020). J Synchrotron Rad 27, 1707–1719.

Gonzalez-Tendero, A., Zhang, C., Balicevic, V., Cárdenes, R., Loncaric, S., Butakoff, C., Paun, B., Bonnin, A., Garcia-Cañadilla, P., Muñoz-Moreno, E., Gratacós, E., Crispi, F. & Bijnens, B. (2017). European Heart Journal - Cardiovascular Imaging 18, 732–741.

Greenspan, H., Van Ginneken, B. & Summers, R. M. (2016). IEEE Trans. Med. Imaging 35, 1153–1159.

Hesamian, M. H., Jia, W., He, X. & Kennedy, P. (2019). J Digit Imaging 32, 582–596.

Kaneko, Y., Shinohara, G., Hoshino, M., Morishita, H., Morita, K., Oshima, Y., Takahashi, M., Yagi, N., Okita, Y. & Tsukube, T. (2017). Pediatr Cardiol 38, 390–393.

Keyriläinen, J., Fernández, M., Fiedler, S., Bravin, A., Karjalainen-Lindsberg, M.-L., Virkkunen, P., Elo, E.-M., Tenhunen, M., Suortti, P. & Thomlinson, W. (2005). European Journal of Radiology 53, 226–237.

Lang, T., Saeidnezhad, N., Dremel, K., Weller, D., Diez, M., Stock, A. M., Sauer, T., Cianciosi, F., Jarnias, C., Tafforeau, P. & Zabler, S. (2023). eJNDT 28, 10.58286/27746.

Lewis, R. A. (2004). Phys. Med. Biol. 49, 3573.

Longo, E., Sancey, L., Cedola, A., Barbier, E. L., Bravin, A., Brun, F., Bukreeva, I., Fratini, M., Massimi, L., Greving, I., Le Duc, G., Tillement, O., De La Rochefoucauld, O. & Zeitoun, P. (2021). Frontiers in Oncology 11.

Lovric, G., Barré, S. F., Schittny, J. C., Roth-Kleiner, M., Stampanoni, M. & Mokso, R. (2013). J Appl Cryst 46, 856–860.

Lyckegaard, A., Johnson, G. & Tafforeau, P. (2011). Int. J. Tomogr. Stat. 18, 1–9.

Madi, K., Staines, K. A., Bay, B. K., Javaheri, B., Geng, H., Bodey, A. J., Cartmell, S., Pitsillides, A. A. & Lee, P. D. (2020). Nat Biomed Eng 4, 343–354.

Massimi, L., Bukreeva, I., Santamaria, G., Fratini, M., Corbelli, A., Brun, F., Fumagalli, S., Maugeri, L., Pacureanu, A., Cloetens, P., Pieroni, N., Fiordaliso, F., Forloni, G., Uccelli, A., Kerlero de Rosbo, N., Balducci, C. & Cedola, A. (2019). NeuroImage 184, 490–495.

Mirea, I., Varray, F., Zhu, Y. M., Fanton, L., Langer, M., Jouk, P. S., Michalowicz, G., Usson, Y. & Magnin, I. E. (2015). Vol. 9126, Functional Imaging and Modeling of the Heart, edited by H. Van Assen, P. Bovendeerd & T. Delhaas. pp. 172–179. Cham: Springer International Publishing.

Mirone, A. (2025). night_rail_bm18 ESRF GitLab.

Mirone, A., Brun, E., Gouillart, E., Tafforeau, P. & Kieffer, J. (2014). Nuclear Instruments and Methods in Physics Research Section B: Beam Interactions with Materials and Atoms 324, 41–48.

Norvik, C., Westöö, C. K., Peruzzi, N., Lovric, G., van der Have, O., Mokso, R., Jeremiasen, I., Brunnström, H., Galambos, C., Bech, M. & Tran-Lundmark, K. (2020). American Journal of Physiology-Lung Cellular and Molecular Physiology 318, L65–L75.

Pacchioni, G. (2019). Nat Rev Phys 1, 100–101.

Paganin, D., Mayo, S. C., Gureyev, T. E., Miller, P. R. & Wilkins, S. W. (2002). Journal of Microscopy 206, 33–40.

Pierantoni, M., Silva Barreto, I., Hammerman, M., Verhoeven, L., Törnquist, E., Novak, V., Mokso, R., Eliasson, P. & Isaksson, H. (2021). Sci Rep 11, 17313.

Pinkert-Leetsch, D., Frohn, J., Ströbel, P., Alves, F., Salditt, T. & Missbach-Guentner, J. (2023). Cancer Imaging 23, 43.

Planinc, I., Garcia-Canadilla, P., Dejea, H., Ilic, I., Guasch, E., Zamora, M., Crispi, F., Stampanoni, M., Milicic, D., Bijnens, B., Bonnin, A. & Cikes, M. (2021). Sci Rep 11, 14020.

Reichardt, M., Frohn, J., Khan, A., Alves, F. & Salditt, T. (2020). Biomed. Opt. Express, BOE 11, 2633–2651.

Reichmann, J., Schnurpfeil, A., Mittelstädt, S., Jensen, P. M., Dahl, V. A., Dahl, A. B., Weide, C., Von Campenhausen, E., Dejea, H., Tafforeau, P., Werlein, C., Jonigk, D., Ackermann, M., Engel, K., Gallwas, J., Dietz, A., Hasanov, M. F. & Salditt, T. (2024). PNAS Nexus 4, pgae583.

Schaeper, J. J., Tafforeau, P., Kampshoff, C. A., Thomas, C., Meyer, A., Stadelmann, C., Liberman, M. C., Moser, T. & Salditt, T. (2025). Npj Imaging 3, 21.

Tapfer, A., Braren, R., Bech, M., Willner, M., Zanette, I., Weitkamp, T., Trajkovic-Arsic, M., Siveke, J. T., Settles, M., Aichler, M., Walch, A. & Pfeiffer, F. (2013). PLoS ONE 8, e58439.

Verleden, S. E., Urban, T., Brunet, J., Wen, W., Peeters, D. J. E., Lapperre, T. S., Walsh, C. L., Tafforeau, P., Dejea, H., Jonigk, D. D., Ackermann, M., Van Hoorenbeeck, K., Lee, P. D., Jacob, J. & Hendriks, J. M. H. (2024). Am. J. Respir. Crit. Care Med. 10.1164/rccm.202403-0509IM.

Vijayakumar, J., Dejea, H., Mirone, A., Muzelle, C., Meyer, J., Jarnias, C., Dollman, K., Zabler, S., Paolasini, L., Bellier, A., Cordonnier, B., Fernandez, V. & Tafforeau, P. (2024). Synchrotron Radiation News 1–10.

Walsh, C. L., Brunet, J., Stansby, D., Gaisné, G., Zhou, Y., Ackermann, M., Bellier, A., Berruyer, C., Bocciarelli, A., Bodin, M., de Bakker, B. S., Dejea, H., De Maria Antolinos, A., Engel, K., Götz, A., Jacob, J., Jonigk, D., Purzycka, J., Urban, T., Verleden, S. E., Xue, R., Tafforeau, P. & Lee, P. D. (2025). bioRxiv 2025.07.31.667856.

Walsh, C. L., Tafforeau, P., Wagner, W. L., Jafree, D. J., Bellier, A., Werlein, C., Kühnel, M. P., Boller, E., Walker-Samuel, S., Robertus, J. L., Long, D. A., Jacob, J., Marussi, S., Brown, E., Holroyd, N., Jonigk, D. D., Ackermann, M. & Lee, P. D. (2021). Nat Methods 18, 1532–1541.

Wang, S., Mirea, I., Varray, F., Liu, W.-Y. & Magnin, I. E. (2019). Vol. Functional Imaging and Modeling of the Heart, edited by Y. Coudière, V. Ozenne, E. Vigmond & N. Zemzemi. pp. 208–216. Cham: Springer International Publishing.

Wilke, J., Krause, F., Niederer, D., Engeroff, T., Nürnberger, F., Vogt, L. & Banzer, W. (2015). Journal of Anatomy 226, 440–446.

Xian, R. P., Brunet, J., Huang, Y., Wagner, W. L., Lee, P. D., Tafforeau, P. & Walsh, C. L. (2024). J. Synchrotron Radiat. 31, 566–577.

Xian, R. P., Walsh, C. L., Verleden, S. E., Wagner, W. L., Bellier, A., Marussi, S., Ackermann, M., Jonigk, D. D., Jacob, J., Lee, P. D. & Tafforeau, P. (2022). Sci Data 9, 264.

Yagis, E., Aslani, S., Jain, Y., Zhou, Y., Rahmani, S., Brunet, J., Bellier, A., Werlein, C., Ackermann, M., Jonigk, D., Tafforeau, P., Lee, P. D. & Walsh, C. L. (2024). Sci Rep 14, 27258.

